# Dynamic *µ*-PBWT: Dynamic Run-length Compressed PBWT for Biobank Scale Data

**DOI:** 10.1101/2025.02.04.636479

**Authors:** Pramesh Shakya, Ahsan Sanaullah, Degui Zhi, Shaojie Zhang

## Abstract

Durbin’s positional Burrows-Wheeler transform (PBWT) supports efficient haplotype matching and queries given a panel of haplotypes. It has been widely used for statistical phasing, imputation and identity-by-descent (IBD) detection. However, the original PBWT panel doesn’t support dynamic updates when haplotypes need to be added or deleted from the panel. Dynamic-PBWT (d-PBWT) solved this problem but it is not memory efficient. While the memory constraint problem of the PBWT has been tackled by Syllable-PBWT and *µ*-PBWT, these are static data structures that do not allow updates. Additionally, Syllable-PBWT only supports long-match query and *µ*-PBWT only supports set-maximal match query, limiting their functionality in the compressed form. In this paper, we present Dynamic *µ*-PBWT (which can also be seen as compressed d-PBWT) that is memory efficient and supports dynamic updates. We run-length compress PBWT to achieve better compression rate and store the runs in the self-balancing trees to enable dynamic updates. We show that the number of updates per insertion or deletion in the tree at each site is constant regardless of the number of haplotypes in the panel and the updates can be made without decompressing the index. In addition, we use orders of magnitude less memory than d-PBWT. We also provide a long match query algorithm that can easily be extended back to the original *µ*-PBWT. Overall, the flexibility and space-efficiency of Dynamic *µ*-PBWT makes it a potential index data structure for biobank scale genetic data analyses. The source code for Dynamic *µ*-PBWT is available at https://github.com/ucfcbb/Dynamic-mu-PBWT.

## 1 Introduction

The positional Burrows Wheeler transform (PBWT) by Durbin [8] supports efficient haplotype matching and queries given a panel of haplotypes. It stores the reverse prefix sorting of the panel of haplotypes at each site and supports haplotype matching within the panel in time linear to the size of the panel. It also supports queries between a query haplotype and the panel in time independent to the size of the panel (only linear to the length of the genome). These efficient query capabilities have been used in many applications such as phasing [3, 7, 12], imputation [16], identity-by-descent (IBD) detection and genealogical analysis [13, 20], and ancestry estimation [21]. For all vs all haplotype matching, memory-efficient scanning algorithms exist. However, for one vs all query searches, the PBWT data structure has to be stored in memory. Hence, the memory efficiency is a bottleneck for fast queries on biobank-scale data.

Several works have been proposed to compress PBWT for efficient storage and to allow fast queries. Syllable-PBWT [19] divides the haplotypes into syllables and builds the PBWT on the compressed syllables reducing the memory usage. It uses a polynomial rolling hash function for string comparison to allow fast long match queries, i.e., identifying all matches longer than a specified length threshold between a query haplotype and the panel. Its compression rate is a function of the syllable size and it offers up to about 100 fold reduction of memory usage. *µ*-PBWT [5] uses run length encoding to compress the PBWT columns, providing a memory efficient alternative for representing the PBWT. It supports set maximal match queries, i.e., all the longest matches between a query haplotype and the panel that are not encompassed by another match. Its compression rate is a function of the number of runs in PBWT columns and offers better compression on panels with sequencing data compared to array data. However, static data structures like PBWT, *µ*-PBWT and Syllable-PBWT do not support adding new haplotypes or removing existing haplotypes from the panel’s PBWT data structure. To update the static PBWT one would have to rebuild it at a cost linear to the size of the panel. This becomes cumbersome to maintain a PBWT data structure for biobank-scale panels for which updates may be frequent (e.g., new participants may be frequently recruited and existing participants may drop off) and rebuilding is expensive as the cost is linear to the size of the panel.

Dynamic PBWT (d-PBWT) [17] solved this problem by replacing the array-based data structures of PBWT with linked lists. d-PBWT provides efficient algorithms to insert and delete a haplotype in time independent of the size of the panel. The authors estimate the time to insert a haplotype into the d-PBWT for the full UK Biobank whole genome array data to be 0.42 seconds and the estimated time to rebuild the PBWT to be 31.25 hours. However, d-PBWT is not memory efficient. It is estimated to consume approximately 29.8 terabytes of memory for holding the UK Biobank (UKB) genotype data [4] with 974,818 haplotypes and 640,000 sites. To make genetic analyses of biobank-scale genotype panel efficient, a data structure that is memory-efficient, supporting frequent haplotype updates, and allowing haplotype matching in its compressed format is needed. The recent trend of moving biobanks to the cloud would further aggravate such needs. For instance, the All of Us Research Program is completely cloud-based [2] and UK Biobank is set to move to a cloud-only environment [18]. This requires analysis tools to be efficient on all aspects as the correlation between computational efficiency and the cost of computation in the cloud is further highlighted with increasing database sizes.

In this paper, we present Dynamic *µ*-PBWT (can also be seen as compressed d-PBWT) that is memory efficient and supports dynamic updates. We bring the flexibility of d-PBWT and the memory efficiencies of the static compressed PBWT together (see Figure 1). To achieve memory efficiency, we run-length compress the PBWT columns and store these run-lengths in self balancing trees to support dynamic updates. This idea of storing run-lengths in self-balancing trees has been used in a BWT context for efficient construction of FM-index for sequencing reads [9, 11]. Here we store the run-lengths of PBWT columns in self balancing trees and show that at each column the number of updates in the tree per insertion and deletion is constant. This makes inserting and deleting a haplotype from the panel efficient. Dynamic *µ*-PBWT consumes 27 to 37 times less memory than d-PBWT but has 11 to 60 times slower insertion time. This is because we do not have constant time access to the haplotype data in the compressed format. Meanwhile, Dynamic *µ*-PBWT has comparable query time with *µ*-PBWT. It consumes approximately 27 to 33 times more memory than *µ*-PBWT but inserts a haplotype in one thousandth of the time it would take *µ*-PBWT to reconstruct with the new haplotype. Overall, dynamic *µ*-PBWT offers a balanced efficiency for biobank-scale genetic analyses.

**Fig. 1.**
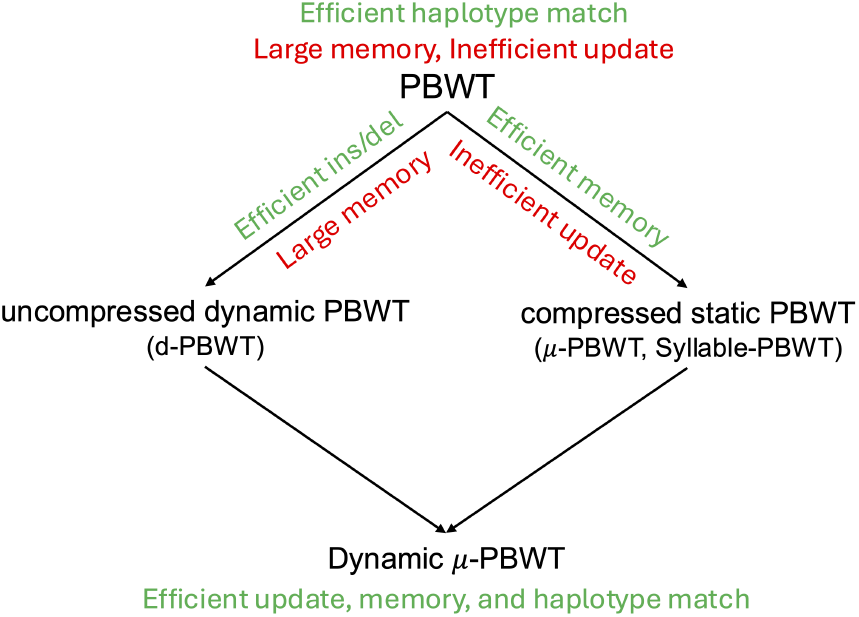
Relationship between PBWT, d-PBWT, compressed static PBWT and Dynamic *µ*-PBWT.

## 2 Preliminaries

### 2.1 Positional Burrows Wheeler Transform

Positional Burrows-Wheeler Transform (PBWT) is a data structure by Durbin that allows efficient haplotype matching and querying [8]. The data structure utilizes a bi-allelic haplotype panel *X* that consists of *M* haplotypes and *N* sites. Each haplotype *x*_*i*_ ∈*X*, 0 ≤*i < M* can have two values at every site i.e. *x*_*i*_[*k*] ∈{0, 1} where 0 ≤*k < N*. We denote the sequence *x*_*i*_ as *x*_*i*_[0, *N*) and its subsequence as *x*_*i*_[*s, e*) where 0 ≤*s < e*≤ *N*. This data structure makes efficient query and haplotype matches by sorting every column of the panel in reverse-prefix order. This sorting groups haplotypes with similar reverse-prefixes. It mainly utilizes two data structures at each column namely, positional-prefix array, *a*_*k*_, and divergence array, *d*_*k*_ for 0≤ *k* ≤*N*. The positional-prefix array, *a*_*k*_, is the permutation of the haplotype indices as a result of co-lexicographic ordering of the prefixes of length *k*. Similarly, the divergence array, *d*_*k*_, stores the starting position of the longest common suffix between a pair of haplotypes in the prefix array. More precisely, *d*_*k*_[*i*] stores the starting position of the longest common suffix between the haplotype corresponding to *a*_*k*_[*i*] and the haplotype in the preceding row in *a*_*k*_[*i* − 1]. A match between two haplotype sequences exists if all the allele values match for a range of sites. A match exists between the haplotype sequence *x*_*i*_ and *x*_*j*_, *i* ≠ *j* if *x*_*i*_[*s, e*) = *x*_*j*_[*s, e*), where 0 ≤ *s < e* ≤ *N* and *x*_*i*_[*s* − 1] *x*_*j*_[*s* − 1] or *s* = 0, and *x*_*i*_[*e*] ≠ *x*_*j*_[*e*] or *e* = *N*. Similarly, a long match is defined as a match that is at least length *L*, i.e. *x*_*i*_[*s, e*) = *x*_*j*_[*s, e*) and *e* − *s* ≥*L*. One of the reasons PBWT is efficient is that its data structures can be constructed for every column in a single scan of the haplotype panel. For a column *k, u*_*k*_ and *v*_*k*_ are arrays of length *M* where *u*_*k*_[*i*] stores the number of zeros before *i* for *i < M* and, *v*_*k*_[*i*] stores the number of ones before *i* for *i < M*. *c*_*k*_ stores the total number of zeros in column *k*. This enables PBWT to map the position of the *i*^*th*^ haplotype from the *k*^*th*^ column to the *k* + 1^*th*^ column with a simple FL mapping, *FL*_*b*_(*i, k*) where *b* is the *i*^*th*^ value in column *k* of PBWT. Therefore, *FL*_*b*_(*i, k*) = *u*_*k*_[*i*] if *b* = 0 and *FL*_*b*_(*i, k*) = *c*_*k*_ + *v*_*k*_[*i*] if *b* = 1.

### 2.2 *µ*-PBWT overview

*µ*-PBWT is a compressed representation of PBWT where every column of PBWT is run-length compressed [5]. A column *k* of PBWT is defined as the *k*^*th*^ column of the haplotype panel where the haplotypes are reverse prefix sorted until *k* − 1. Since every column of PBWT is run-length compressed, *r*_*k*_ denotes the number of runs in column *k* of the PBWT, 0≤ *k < N*, and *r* denotes the total number of runs in the PBWT panel, 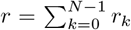. Each column of PBWT is summarized with a few data structures. The following are the *µ*-PBWT data structures for the *k*^*th*^ column of PBWT: *p*_*k*_[*i*] is head of a run i.e. the first index in the *i*-th run of the *k*^*th*^ column, *c*_*k*_ stores the number of zeros in column *k, b*_*k*_ stores the bit-value of the first run in column *<u>k</u>* and *uv*_*k*_[*i*] is the number of *b*_*k*_ bits before the *i*^*th*^ run in column *k* if *i* is odd, otherwise it is the number of 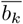 bits before the *i*^*th*^ run in column *k*.

The above data structures called the mapping structure allow us to move from *k*^*th*^ column to *k* + 1^*th*^ column in *µ*-PBWT. It should be noted that *µ*-PBWT only stores prefix array values at the beginning and end of each run at each column and stores divergence values at the beginning of each run at each column.

It also stores thresholds at each column which gives the position of the first maximum divergence value (minimum divergence value if divergence value is defined as length as in *µ*-PBWT) in each run. Hence, the authors describe a *ϕ*(*i, k*) function which returns the value preceding *i* in *a*_*k*_. i.e. if *a*_*k*_[*j*] = *i, ϕ*(*i, k*) = *a*_*k*_[*j* − 1]. Its inverse, *ϕ*^−1^(*i, k*) returns the value following *i* in *a*_*k*_, therefore *ϕ*^−1^(*i, k*) = *a*_*k*_[*j* + 1].

In this paper, we define a function *D*_*k*_(*i*) that returns the divergence value of haplotype *i* in column *k* where *a*_*k*_[*j*] = *i*. By definition, this is the starting position of the match between *i* and *a*_*k*_[*j* − 1]. If *i* is not at the beginning of a run in column *k*, we wouldn’t have access to the divergence value. Since divergence values are only sampled at the beginning of each run, we take the first column after *k* (inclusive) where haplotype *i* is at the top of a run. Say, *j*, 0 ≤*k* ≤*j < N*, is the first column after *k* where haplotype *i* is at the top of a run and *i* maps to row *i*^*′*^ in *a*_*j*_, *a*_*j*_[*i*^*′*^] = *i*, then the divergence value sampled here would be the same for haplotype *i* at column *k*. Therefore, *D*_*k*_(*i*) = *d*_*j*_[*i*^*′*^]. It should be noted that *µ*-PBWT defines divergence value as the length of the match which is functionally equivalent to our definition as the starting position of the match. *D*_*k*_(*i*) can be computed efficiently by having an array for each haplotype when it is at the top of a run. Then a binary search can be performed to find the column after *k* (including) when *i* is at the top of the run. An efficient implementation can use any predecessor/successor data structure, for example, a sparse-bit vector.

## 3 Methods

### 3.1 Dynamic *µ*-PBWT

The Dynamic *µ*-PBWT transforms the *µ*-PBWT into a dynamic compressed structure. First, we run-length compress the PBWT columns inspired by *µ*-PBWT. Then, we make it dynamic by storing runs in a self-balancing tree in each column. Typically, dynamic bit vectors store the sequence of bits in the leaves of a self balancing tree [14, 15]. However, in our case, we propose to store run-length compressed bits in the leaves of such trees. In theory, any self-balancing tree could be used for this purpose but we choose B+ tree for memory efficiency and better cache usage. The key idea here is to maintain a B+ tree at each column of PBWT to store the runs in its leaves to allow efficient insertion and deletion of a haplotype.

B+ trees are generalized binary search trees where each node can store multiple keys. A B+ tree consists of a root node, internal nodes and leaf nodes. Note that B+ trees only store data at the leaf nodes unlike B trees. It is defined by the minimum degree *t, t*≥ 2, where each node can have at most 2*t*− 1 keys and each node should have at least *t* −1 keys (except the root node). A node is considered full when it has the maximum number of keys. Each node *x* stores *x.n* keys and have *x.n* + 1 children. Each key in an internal node stores information about the number of bits in *x.key*_*i*_.*num*, number of ones in *x.key*_*i*_.*ones* and number of runs in *x.key*_*i*_.*runs* for 1 ≤*i* ≤*x.n*. We define the number of bits in a tree rooted at *y* as the sum of the number of bits in all of its subtrees. Then, *x.key*_*i*_.*num* stores the sum of the number of bits in all the subtrees to the left of it i.e. sum of the number of bits in all the subtrees rooted at *x.c*_*k*_ where 1 ≤*k* ≤*i*. Similarly, *x.key*_*i*_.*ones* stores the number of ones in all the subtrees rooted at *x.c*_*k*_ for 1 ≤*k* ≤*i* and *x.key*_*i*_.*runs* store the number of runs in all the subtrees rooted at *x.c*_*k*_ for 1≤ *k*≤ *i*. However, the keys in a leaf node store a pair of runs. The *i*^*th*^ key in a leaf node stores the number of bits in a run of zeros in *x.key*_*i*_.*f* and the number of bits in a run of ones in *x.key*_*i*_.*s* in this order. Here onwards, we use trees to refer to B+ trees for simplicity.

Figure 2 shows the tree representation of a run-length compressed PBWT column. Figure 2A shows a haplotype panel of 20 haplotypes and 15 sites. This panel is reverse prefix sorted by the reversed prefixes of length *k* for each column *k* and those PBWT columns are shown in Figure 2B. The light gray squares show the runs of zeros and the dark gray squares show the run of ones. Each PBWT column is then run-length compressed to obtain Figure 2C. Figure 2D shows the B+ tree of minimum degree *t* = 2 storing the runs of the second column of PBWT, i.e., 00001000010010110000. We observe that there are *r*_1_ = 9 runs in this PBWT column. Each key of the leaf stores a pair of runs distinguished by the light and dark gray bands below the leaves. The leftmost key of the root node shows that there are *x.key*_1_.*num* = 10 bits in leaves of all the subtrees to the left of it. Of the 10 bits *x.key*_1_.*ones* = 2 are ones and they all together constitute *x.key*_1_.*runs* = 4 runs. Note that the root’s second key stores information about the runs in all the child subtrees to the left of it. Hence, *x.key*_2_.*num* = 16 as it stores the number of bits in the two leaves to the left of it. Similarly, *x.key*_2_.*ones* = 5, and *x.key*_2_.*runs* = 8. The leftmost key in the leftmost leaf node stores the run of zeros with *x.key*_1_.*f* = 4 bits and the run of ones with *x.key*_1_.*s* = 1 bits. The rightmost key of the rightmost leaf node stores only the run of zeros as *x.key*_1_.*f* = 4 and *x.key*_1_.*s* = 0.

**Fig. 2.**
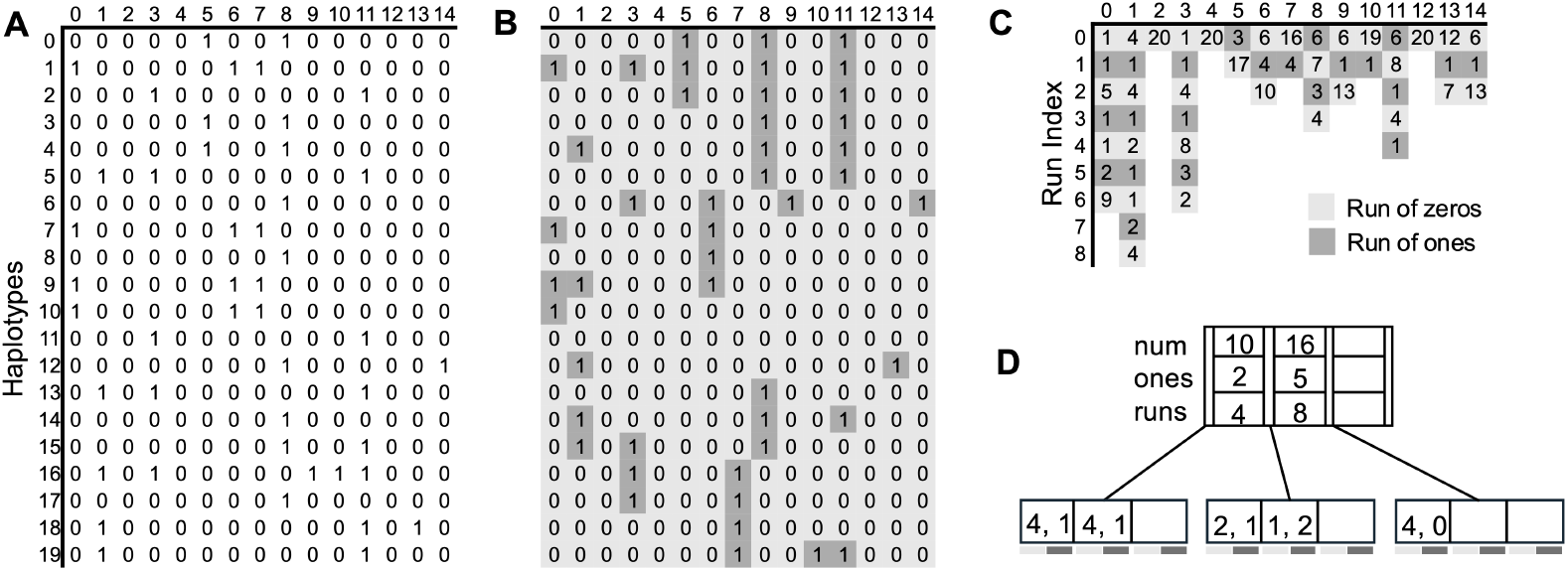
Dynamic run-length compressed representation of the PBWT panel of haplotypes. (A) A bi-allelic panel consisting of 20 haplotypes (rows) and 15 sites (columns). (B) PBWT of the input panel of haplotypes. Light gray and dark gray squares show a run of zeros and ones respectively. (C) Run-length encoded representation of PBWT columns. (D) B+ tree (*t* = 2) representation of the second column of PBWT. The light and dark bands below the leaves show runs of zero and ones respectively.

#### FL mapping

We require the primary functionality in PBWT of mapping *i*^*th*^ haplotype in column *k* to column *k* + 1. We need *u*_*k*_[*i*] and *v*_*k*_[*i*] values for this. We compute this by traversing the tree at column *k* from the root node to the leaf node tracking the number of zeros and number of ones that exist before *i*. Then, we find the respective key in the leaf node and calculate the *u* and *v* values from the run lengths stored at the leaf nodes. We define *uv*(*x, i, z, o*) as a function that calculates the *u* and *v* values before *i* in the tree rooted at *x*. We track the number of zeros and ones we have seen so far with *z* and *o*. For a given index *i*, we start at the root node and iterate through the keys to find the first key with *num* greater than *i*. If *x.key*_*j*_ is the first key such that *x.key*_*j*_.*num > i* and 1 ≤ *j* ≤ *x.n*, we traverse down the corresponding child node *x.c*_*j*_ with adjusted index *i* = *i*− *x.key*_*j*−1_.*num, j >* 1. The number of zeros and ones seen so far is calculated from the preceding key as *z* = *x.key*_*j*−1_.*num*− *x.key*_*j*−1_.*ones* and *o* = *x.key*_*j*−1_.*ones*. When we are at the leaf node, we find the key and the run within the key where *i* falls into. We do this by iterating over all the keys tracking the cumulative sum of the values of the keys in *sum*. The first key where the sum is greater than *i* is our desired key. Next, we find if *i* is in a run of zeros or a run of ones within the key. It is in a run of ones if the index is greater than or equal to the difference of the cumulative sum and the number of bits in the run of ones, i.e., *i*≥ *sum*− *x.key*_*i*_.*s*. If not, it lies in the run of zeros. We can calculate the offset of *i* from the head of run and get the number of zeros and ones before *i* (see Algorithm 1 in Appendix). The time complexity to compute this is *O*(log *r*_*k*_). This enables us to successfully map the *i*^*th*^ haplotype from column *k* to column *k* + 1. We define this mapping functionality as an extension function *w*(*x, i, k, v, id*), *v*∈{0, 1} where it maps haplotype *id* at the *i*^*th*^ position in column *k* with allele *v* to column *k* + 1 and also returns the haplotype id in column *k* + 1. *x* is the root node of the tree at column *k* (see Algorithm 2 in Appendix). Note that we can also find the run index of *i*^*th*^ haplotype at a column by traversing the tree in a similar manner.

#### B+ trees for Dynamic *µ*-PBWT

For each column of PBWT, we store the run lengths, prefix array samples at the start and end of each run, and the divergence values at the start of each run in B+ trees for efficient insertion and deletion of a haplotype in Dynamic *µ*-PBWT. For each haplotype, we also maintain a B+ tree that stores the haplotypes above it for all the columns where it is at a start of a run. Similarly, for each haplotype, we maintain another B+ tree that stores the haplotypes below it for all the columns where it is at the bottom of a run. These haplotypes are stored in the B+ tree in order of the columns where they appear at the start or bottom of a run. This allows us to efficiently update the tree when a new haplotype is inserted in place of the other haplotype at either of the run boundaries. This enables the *ϕ* functionality from *µ*-PBWT in Dynamic *µ*-PBWT. Additionally, we also store the total number of zeros in each column and the starting bit at each column.

There are some key differences in the way we update the B+ trees in Dynamic *µ*-PBWT from general B+ trees. During insertion, we traverse the tree from the root node to the leaf node and find the right key of the leaf node to update. With each insertion, the appropriate keys (and their parameters) of the internal nodes in the path are also updated. Whenever a node is full, i.e. there are 2*t* − 1 keys, the node is split into two nodes. First, we explain how to split a full node *x.c*_*i*_ with a non full parent node *x*. The splitting of a leaf node is handled differently from that of an internal node. When splitting a leaf node *x.c*_*i*_, first we create a new leaf node. Then, *t* − 1 keys after the median key in *x.c*_*i*_ are moved to the new leaf node. The split leaf node *x.c*_*i*_ will contain *t* keys instead of *t* − 1 keys. A new key is inserted in the parent node *x* as *x.key*_*i*_ where *x.key*_*i*_.*num* is the number of bits in *x.c*_*i*_, *x.key*_*i*_.*ones* is the number of ones in *x.c*_*i*_ and *x.key*_*i*_.*runs* is the number of runs in *x.c*_*i*_. Finally, all the keys following *x.key*_*i*_ (including *x.key*_*i*_) in *x* are updated to reflect the changes after insertion (see Algorithm 3 in Appendix).

If the parent node *x* is not full after inserting the new key *x.key*_*i*_, we stop. However, if it is full, we split it following the rules of splitting an internal node. Here, we define *x.c*_*i*_ as the full internal node and *x* as its non full parent node. To split *x.c*_*i*_, first we create a new internal node. The *t* − 1 keys to the right of the median key of *x.c*_*i*_ are moved into the new node. For every key moved into the new node, its parameters are adjusted by subtracting its *num, ones*, and *runs* values from its preceding key. Then the median key is inserted into the parent node as *x.key*_*i*_ and all the keys following *x.key*_*i*_ (including *x.key*_*i*_) are updated (see Algorithm 4 in Appendix). As in B+ tree, the split is propagated upwards until a non-full internal node is found. If we do not find a non-full internal node and the split continues to the root node, a new root node as the parent of the existing root node is created. Then, the old root node is split. This would increase the height of the tree by one.

### 3.2 Insertion

#### Update insert positions and divergence values in PBWT

The insertion algorithm inserts a new haplotype *z* into the Dynamic *µ*-PBWT. First, the insert positions in PBWT of the new haplotype are calculated at each column. We compute this by tracking the position of the haplotype that would be below the inserted haplotype at each column if *z* were in the PBWT. We define *t*_*k*_ as the haplotype that would be below the inserted haplotype at column *k. t*_*k*_ stores two parameters *t*_*k*_.*pos* and *t*_*k*_.*ID. t*_*k*_.*pos* is the index of haplotype *t*_*k*_.*ID* in *a*_*k*_, i.e., *a*_*k*_[*t*_*k*_.*pos*] = *t*_*k*_.*ID*. We map the position of this haplotype from column *k* to column *k* + 1 using the extension function as *t*_*k*+1_ = *w*(*x*_*k*_, *t*_*k*_.*pos, k, z*[*k*], *t*_*k*_.*ID*). This process is the same as computing the matching statistic of *µ*-PBWT and the virtual insertion algorithm in d-PBWT.

The next step is to update the divergence values of the inserted haplotype, *z*, and the haplotype below it at every column. To compute this, we scan the panel from right to left and compute the arrays *zStart* and *tStart*. We define *zStart* and *tStart* as two arrays of length *N* + 1 that store the divergence values of *z* and *t*_*k*_ at each column if *z* were in the PBWT. *zStart*[*k*] is the divergence value of *z* at column *k* and *tStart*[*k*] is the divergence value of *t*_*k*_ at column *k*. Therefore, for 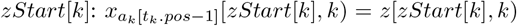 where 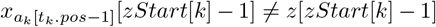 or *k* = 0 and, 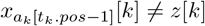 or *k* = *N*. Similarly, for 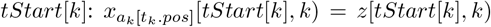 where 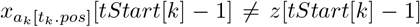 or *k* = 0 and 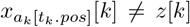 or *k* = *N*. For haplotype *z*, we need to access the haplotype above it to compute the divergence values. We get this haplotype using *ϕ*(*t*_*k*_.*ID, k*). We define *beg*_*h*_ as the number of times haplotype *h* is at the beginning of a run across all columns and, *end*_*h*_ as the number of times haplotype *h* is at the end of a run across all columns. Then, the time complexity to compute *ϕ*(*h, k*) is *O*(log *beg*_*h*_). Similarly, the time complexity to compute *ϕ*^−1^(*h, k*) is *O*(log *end*_*h*_). Next, we compare the alleles between *z* and *ϕ*(*t*_*k*_.*ID, k*) until there’s a mismatch. Similarly, the divergence value for *t*_*k*_ is also computed by comparing the alleles between *t*_*k*_.*ID* and *z* until there’s a mismatch (see Algorithm 5 in Appendix). It should be noted that the divergence value at any column *k* for any haplotype is at least the divergence value at *k* + 1 column, i.e., *d*_*k*_[*i*] ≤ *d*_*k*+1_[*i*] [17]. Therefore, the divergence values should be updated in *O*(*N*) runtime if we have constant time access to any haplotype. However, we do not have constant time access to this information as we only store prefix array samples at the start and end of each run at each column and only store the divergence values at the start of each run at each column. Hence, the computation of divergence values is worst case *O*(*N*^2^ log *a* + *N* log *b*). Here, *a* is the maximum number of runs across all columns, i.e., *a* = max(*r*_*k*_), 0 ≤ *k < N* and, *b* is the maximum number of times any haplotype is at the start or end of a run, i.e., *b* = max(max(*beg*_*h*_), max(*end*_*h*_)) for 0 ≤ *h < M*.

#### Update B+ trees

The final step is to update all Dynamic *µ*-PBWT B+ trees one column at a time in a forward sweep. Here, we explain the process of updating the runs stored in a B+ tree at a single column. This process will be the same to update all other B+ trees of Dynamic *µ*-PBWT.

The insertion in a column can result in three scenarios, namely, updating the existing run, inserting a new run or splitting the existing run to insert a new run. We define the insertion operation at a column as *Insert*0(*x, i*) when we’re inserting 0 at index *i* in a tree rooted at *x*. Similarly, we define *Insert*1(*x, i*) as insertion of 1 at index *i* in a tree rooted at *x*. The simplest case is when the inserted bit matches the bit-value of the run at the insert location. Here, the existing run is incremented by 1. The equivalent tree operation is to traverse the tree down to the leaf node to increment the value in the corresponding key of the leaf node by 1. Note that all the keys and their parameters are updated to reflect the change on the path from the leaf node back to the root node (see Algorithms 6 and 7 in Appendix).

The second case of inserting a new run only happens when inserting at the top or bottom of a column and the inserted bit does not match the bit-value of the run. The equivalent tree operation would be to insert a new key in the leftmost or rightmost leaf node. We traverse the tree from the root node to the leaf node and insert the new run as a key. If the leaf node becomes full, it is split and the split is propagated upwards to the root. A special case is inserting a new run of one at the top of a column that starts with a run of zeros. This is equivalent to inserting a key with value [0, 1] in the left-most leaf node. We traverse the tree down to the leftmost leaf node. When we’re at the leftmost leaf node, we shift all the existing keys to the right by one position and insert the new key at the leftmost position as *x.key*_1_.*f* = 0 and *x.key*_1_.*s* = 1. If the leaf node becomes full, we split it as discussed before. Note that, if we’re inserting at run boundaries (except when it’s at the top or bottom of a column) and the inserted bit does not match the bit-value of the run, the neighboring run is updated.

Finally, the third case of a run being split happens when we insert a bit that does not match the bit-value of the run and it is not at a run boundary. For example, if we insert a 1 in a run of zeros or vice-versa, the existing run would be split to insert the bit. This is equivalent to inserting the new key in the leaf node. If *x.key*_*j*_ contains the run to be split, we move all the keys from *j* + 1 to *x.n* to the right by one position. We then insert a new key in *j* + 1^*th*^ position. If a run of zeros is being split, we move the run of ones at *x.key*_*j*_.*s* to the new key i.e., *x.key*_*j*+1_.*s* = *x.key*_*j*_.*s*. Then, we calculate the left half of the split and the right half of the split. The left half of the split is assigned to *x.key*_*j*_.*f*. The new inserted bit is updated at *x.key*_*j*_.*s* = 1. Then, the right half of the split is updated at *x.key*_*j*+1_.*f*. If inserting this new key makes the leaf node full, it is split and the split is propagated upwards (see Algorithm 8 in the Appendix). We observe that for each of these cases, the number of updates made in the tree is constant however, the time complexity for each of these updates is a function of the number of runs in the tree.

Figure 3 shows the different cases when a new haplotype *z* is inserted. Figure 3A shows the run-length compressed panel before and after insertion. The black boxes show the insert positions of the new haplotype *z* in each column. The figure on the right shows the updated panel after insertion. The existing runs being updated after insertion are shown by the orange boxes. A new run is being inserted at the bottom of column 7 as the inserted bit 1 does not match the bit-value of the last run i.e. 0. The new runs created are shown by blue boxes. Columns 3 and 6 show the runs being split after inserting a 1 in a run of zeros. The run that was split is shown by a dotted orange box. Note that every time a run is split, the total number of runs in a column increases by two. Figure 3B shows how the tree corresponding to column 3 is updated during insertion. The bold letters show the values that are to be updated or have been updated. The leftmost tree is the initial tree where the leaf node with the pair of values [8, 3] is about to be updated. Since the run of eight zeros is being split, a new key [4, 3] is inserted into the leaf node as shown in the middle tree. As this leads to the leaf node to become full, we split the leaf node as shown in the rightmost tree. Here, a new key is added to the root node (shown by the bold letters) which show the number of bits, number of ones and number of runs in the leaves of all the subtrees to the left of it.

**Fig. 3.**
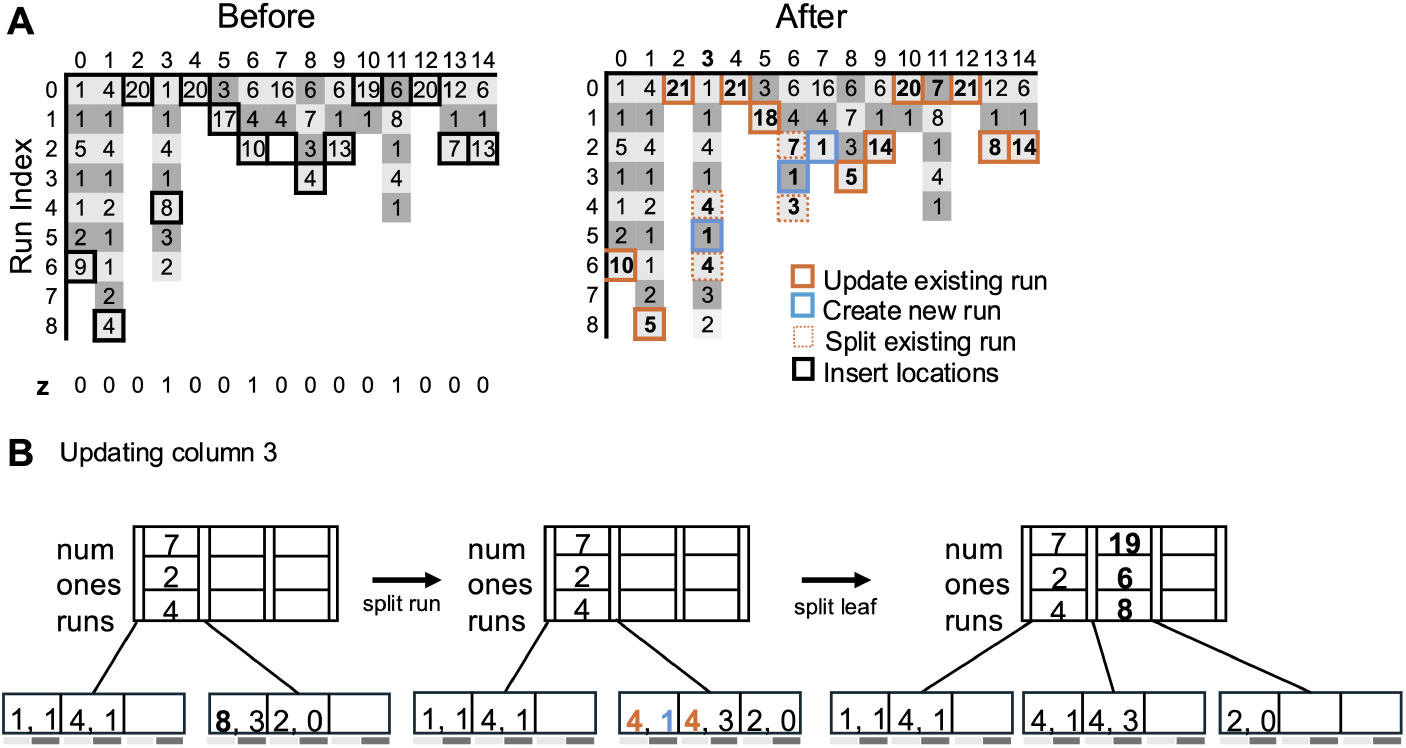
Insertion of a new haplotype in Dynamic *µ*-PBWT. (A) Dynamic *µ*-PBWT run lengths before and after insertion of the new haplotype. New haplotype *z* is shown at the bottom of the left figure. The haplotype’s insert locations are shown by bold black boxes. The right figure shows the panel after insertion. Orange boxes show the existing runs that were updated after insertion. Blue boxes show new runs being inserted and dotted orange boxes show the existing run being split to insert a new run. (B) The corresponding B+ tree operation when updating column 3.

So far, we explained the process of updating the runs stored in a B+ tree. Since we also store the prefix array samples and divergence value samples in B+ trees, these are also updated during insertion. The prefix array samples are only updated when the new haplotype is inserted at the start or end of a run. This is equivalent to updating the leaf of a tree to replace the old start or end of a run with the new haplotype. The prefix array samples are also updated when an existing run is split. In this case, the inserted haplotype and the haplotype below it become the new start of runs. Hence, these haplotypes are inserted into the tree. Similarly, the divergence values are also updated when the new haplotype is inserted at the start or the end of a run. If the new haplotype is inserted at the start of a run, the divergence value of the old start of run is replaced by the divergence value of the newly inserted haplotype (except when it is at the top of a column). However, if the new haplotype is inserted at the end of a run, the divergence value of the haplotype below is updated (except when it is at the bottom of the column). The *ϕ* data structure is also updated to store a B+ tree for the newly inserted haplotype to contain all the haplotypes above it and below it for any column where it is at the start or end of a run. Additionally, for any haplotype when the haplotypes above and below change at the run boundaries because of the inserted haplotype, those are also updated accordingly.

The time complexity to update the trees across all columns is *O*(*N* log *a* + *N* log *b*). Hence, the overall time complexity of insertion is *O*(*N*^2^ log *r*_*max*_), where *r*_*max*_ = max(*a, b*).

### 3.3 Deletion

To delete a haplotype from Dynamic *µ*-PBWT, first we find the positions of the haplotype to be deleted in PBWT and its respective bit values in all the columns. This can easily be computed using the extension function. Then we delete a single bit at each position one column at a time. We define *Delete*0(*x, i*) as deletion of a 0 bit at index *i* in a tree rooted at *x* in a column. Similarly, we define *Delete*1(*x, i*) as deletion of 1 at index *i* in a tree rooted at *x* in a column. We delete a bit by traversing the tree from the root node to the leaf node as before. When we are at the leaf node, we locate the right key and decrement the run value by one. The special case of deletion occurs when we delete a run. In this case, if we delete a run of ones, then the two flanking runs of zeros will be merged and vice-versa (except when the deleted run is at the top or bottom of a column). If a run of zeros is deleted from a leaf node *x* at the key index *i*, this would result in *x.key*_*i*_.*f* = 0. Then, we would merge the run of ones of the current key with a predecessor key within the same node. If the predecessor key within the same node does not exist we merge it with the last key of a predecessor leaf node. Similarly, if a run of ones would be deleted, the current key’s run of zeros is merged with a successor key in the same node or the first key in the successor leaf node. After merging the values of the current key and updating the internal nodes’ keys (and their parameters) appropriately we can delete the key from the tree following B+ tree deletion. If leaf node has less than the minimum number of keys after deletion, the tree will be rebalanced to maintain B+ tree properties (see Algorithms 9 and 10 in Appendix). Similar to the insertion process, we also update the prefix array samples, divergence samples and the *ϕ* data structure. The overall time complexity for deletion operation is *O*(*N* log *r*_*max*_).

### 3.4 Long match query

The haplotype long match query problem is to find all matches that are at least length *L* between a query haplotype *z* and the Dynamic *µ*-PBWT (or static *µ*-PBWT) index of haplotype panel X. To find all long matches, we virtually insert the query haplotype, compute the divergence values and find the matches that satisfy the length cutoff.

We virtually insert the query haplotype *z* into the panel by tracking the haplotype that would be below *z* if it was inserted into the panel. Then, we calculate the divergence values for the query haplotype and the haplotype below it if *z* would be in the panel. This process is similar to the insertion process described in section 3.2. Once the divergence values are updated, the *L*-long matches are found by tracking the matching blocks. Matching blocks are the block of sequences that match with *z* at least length *L*. Here, we adapt the long match query algorithm from d-PBWT [17] to Dynamic *µ*-PBWT data structures. Algorithm 11 describes the details of using Dynamic *µ*-PBWT data structures. We use *D*_*k*_(*i*) to access the divergence value for haplotype *i, a*_*k*_[*j*] = *i*, and *ϕ*(*i, k*) (*ϕ*^−1^(*i, k*)) to access the haplotype preceding (or following) haplotype *i* in column *k* (see Algorithm 11 in Appendix). The time complexity of virtually inserting the haplotype and calculating the divergence values is *O*(*N*^2^ log *a* + *N* log *b*) where *a* is the maximum number of runs across all columns and *b* is the maximum number of times any haplotype is at the top or bottom of a run. The time complexity of the query algorithm is *O*(*output* log *r*_*max*_), where |*output*| is the total number of matches outputted and *r*_*max*_ = max(*a, b*). Hence, the overall long match query algorithm is *O*(*N*^2^ log *r*_*max*_ + |*output*| log *r*_*max*_). It should be noted that the process of virtual insertion and divergence value calculation is similar to the computation of the matching statistic data structure, *A*, in *µ*-PBWT [5]. Hence, this algorithm can be easily extended to support long match query in *µ*-PBWT.

## 4 Results

### 4.1 Data

We tested Dynamic *µ*-PBWT on datasets of UK Biobank (UKB) [4] and 1000 Genomes Project (1KGP) [1]. The 1KGP dataset is filtered to retain only bi-allelic SNPs using bcftools [6] with the command “bcftools view -m2 -M2 -vSNPS”. The UKB dataset has 974,818 haplotypes (487,409 individuals) and approximately 700,000 sites. Similarly, the 1KGP dataset has 5008 haplotypes (2504 individuals) and 77,818,346 sites after filtering. We use the DYNAMIC [15] library to implement B+ trees and implement the Dynamic *µ*-PBWT data structures.

### 4.2 Construction time and memory

We compared the construction time and memory consumption of Dynamic *µ*-PBWT with *µ*-PBWT, Syllable-PBWT and d-PBWT. For the comparison on UKB data, we randomly sample 50,000 haplotypes for each chromsome. For 1KGP data, we use 5008 haplotypes across all autosomes. We compared the construction time between Dynamic *µ*-PBWT, *µ*-PBWT and Syllable-PBWT on both datasets. We do not compare construction time with d-PBWT as it does not have a separate construction method. The construction time of Dynamic *µ*-PBWT is comparable to *µ*-PBWT on both datasets whereas Syllable-PBWT is roughly 5-7 times faster than both Dynamic *µ*-PBWT and static *µ*-PBWT as shown in Figure 4.

**Fig. 4.**
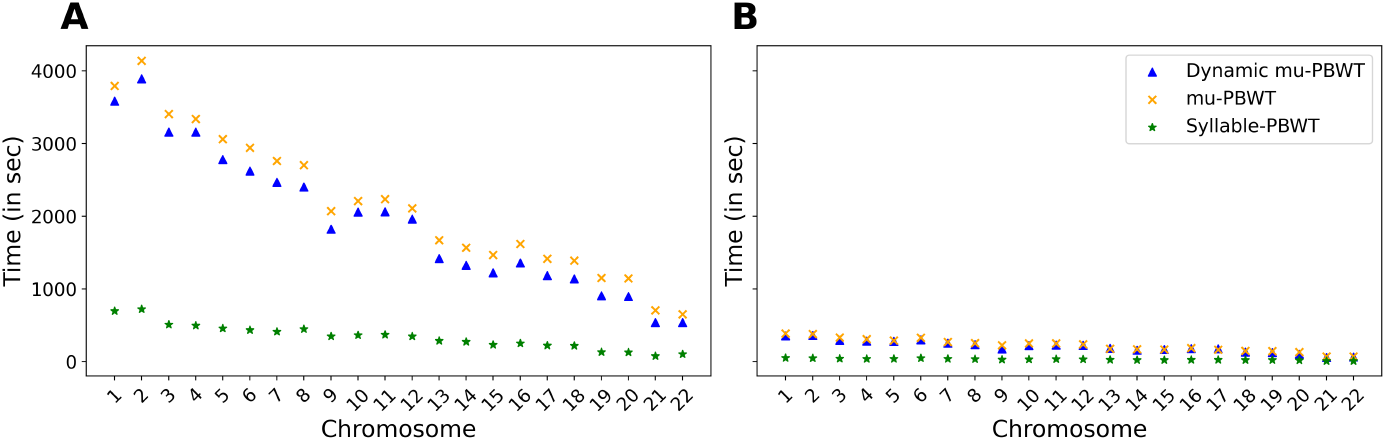
Construction time comparison between Dynamic *µ*-PBWT, *µ*-PBWT and Syllable-PBWT. (A) Construction time comparison on all autosomes of 1KGP data (B) Construction time comparison on all autosomes of UKB data with randomly sampled 50000 haplotypes.

Figure 5A shows the memory comparison on 1KGP data and Figure 5B shows the memory comparison on the subsetted panel of UKB data. The gray + sign in both figures show the estimated d-PBWT memory usage. We estimated d-PBWT memory usage on these data using 48 bytes per site per haplotype estimate from the authors [17]. We observe that Dynamic *µ*-PBWT consumes approximately 26 times less memory than d-PBWT on all the autosomes of 1KGP data but it consumes roughly 27 to 33 times more memory than *µ*-PBWT and Syllable-PBWT. On the subsetted UKB data, Dynamic *µ*-PBWT consumes approximately 2.5 times more memory than *µ*-PBWT and 8-12 times more memory than Syllable-PBWT but it consumes 36 times less memory than d-PBWT. We show that Dynamic *µ*-PBWT consumes more memory than its static compressed counterparts to support dynamic updates. However, it consumes significantly less memory compared to d-PBWT.

**Fig. 5.**
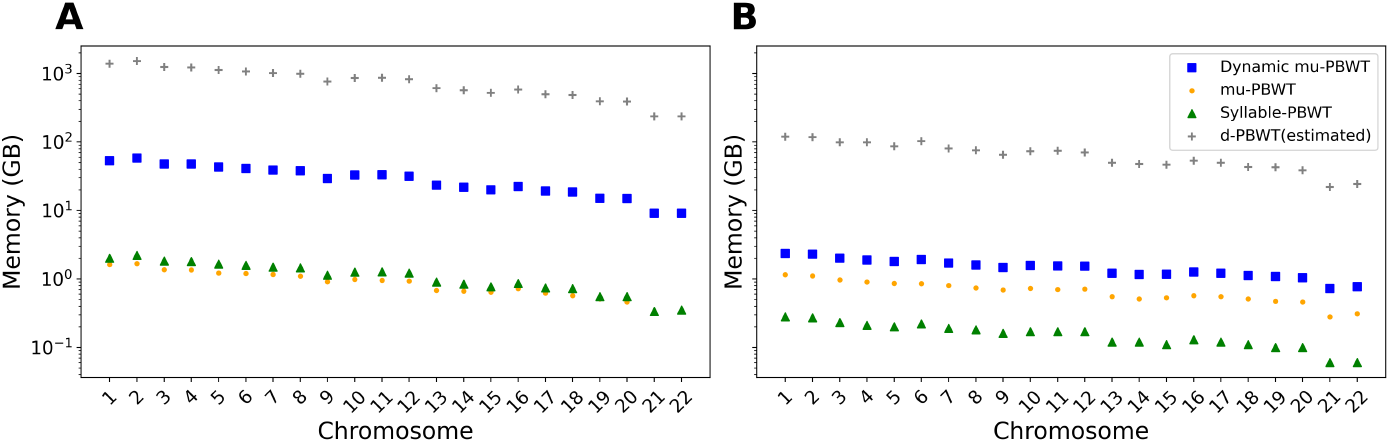
Memory comparison between Dynamic *µ*-PBWT, *µ*-PBWT, Syllable-PBWT and d-PBWT. (A) Memory comparison on all autosomes of 1KGP data (B) Memory comparison on all autosomes of UKB data with randomly sampled 50000 haplotypes.

### 4.3 Insertion and deletion

We tested the insertion and deletion time of Dynamic *µ*-PBWT on chromosome 21 of 1KGP data with 1,054,447 sites and on a subpanel of chromosome 21 of UKB data with 9793 sites. For the UKB data, we randomly sampled 100,000 haplotypes from 974,818 haplotypes. We test the insertion time by inserting one haplotype at a time into the Dynamic *µ*-PBWT in random order. We test the deletion time by first building the Dynamic *µ*-PBWT and deleting one haplotype at a time in random order. Each insertion and deletion was timed. The insertion times and deletion times per haplotype were averaged every 100 haplotypes and those 1000 points are plotted in Figure 6. Figure 6A shows the average insertion time per haplotype for 1KGP data in relation to the database size respectively. Figure 6B shows the average deletion time per haplotype on 1KGP data. Figure 6 (C, D) shows the average insertion and deletion time for UKB data in relation to the database size respectively. We observe that it takes approximately 1500 milliseconds to insert a single haplotype and approximately 3000 milliseconds to delete a single haplotype for chromosome 21 of 1KGP data. For the subsetted chromosome 21 panel of UKB data, it takes approximately 150 milliseconds to insert and approximately 350 milliseconds to delete a single haplotype. The insertion and deletion time increases sublinearly to the number of haplotypes for the UKB data. This is expected as the insertion and deletion times are a function of the number of runs in the PBWT. However, this relationship is as not as distinct for 1KGP data. This may be due to the smaller panel size of 1KGP data. The deletion time is roughly two times that of insertion time because we reinsert the bottom haplotype inplace of the deleted haplotype when the haplotype is deleted from the middle of the panel. We reinsert the bottom haplotype so that the haplotype indices in the panel remain in the range [0, *M*).

**Fig. 6.**
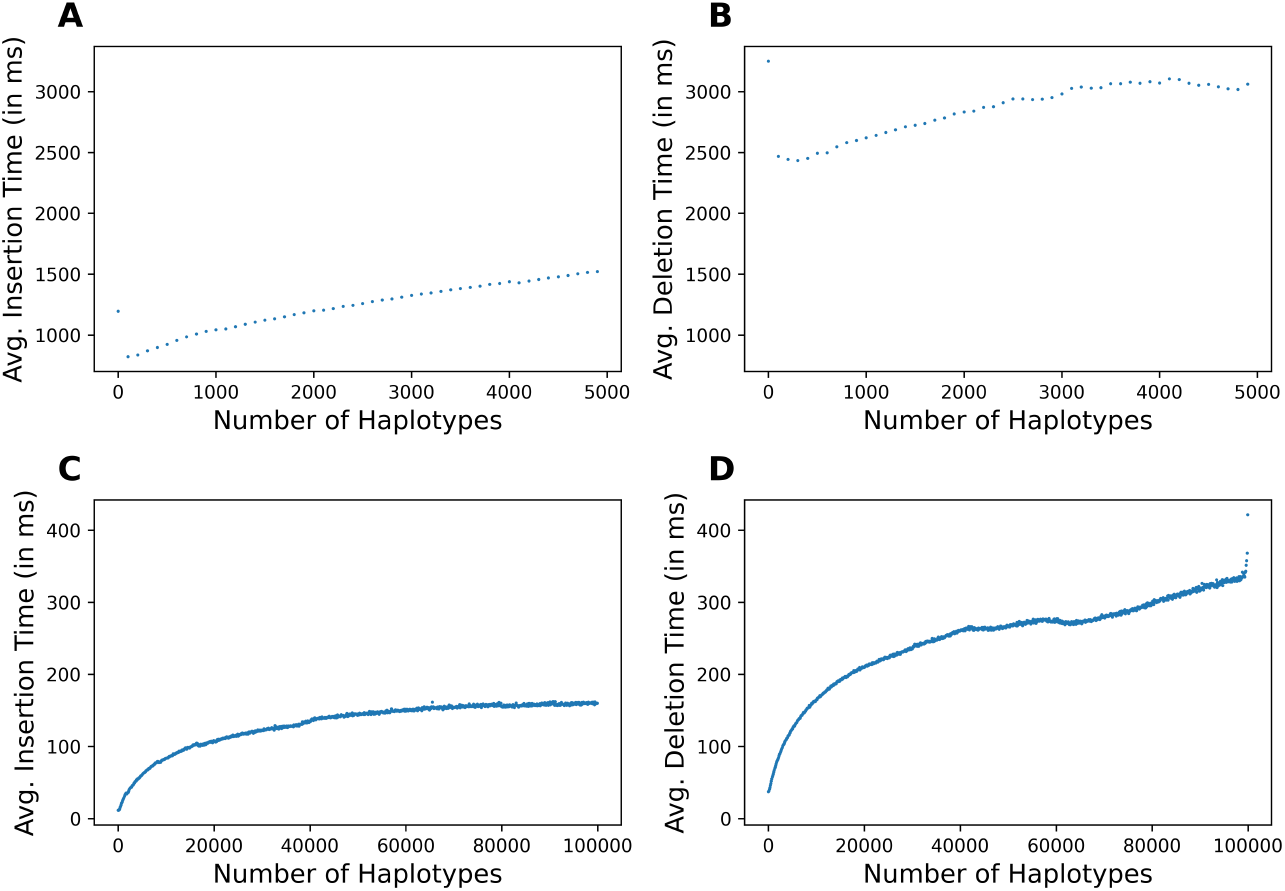
Time for insertion and deletion of haplotypes. (A, B) Average insertion and deletion time of 5008 haplotypes of chromosome 21 of 1KGP. (C, D) Average insertion and deletion time of 100,000 haplotypes of chromosome 21 of UK Biobank data.

The average insertion time of d-PBWT for chromosome 21 of 1KGP was 0.27 seconds and the average deletion time was 0.21 seconds. The average insertion time of d-PBWT on the subpanel of UKB data was 0.0025 seconds and the average deletion time was 0.0022 seconds. Dynamic *µ*-PBWT consumed 9.09 GB and d-PBWT consumed 241.43 GB for 1KGP data. Dynamic *µ*-PBWT consumed 1.31 GB and d-PBWT consumed 44.83 GB on the subpanel of UKB data. For 1KGP data Dynamic *µ*-PBWT is roughly 5-14 times as slow as d-PBWT but takes 27 times less memory than d-PBWT. For the subpanel of UKB data, Dynamic *µ*-PBWT is roughly 60-160 times slower than d-PBWT but consumes approximately 37 times more memory than d-PBWT. Therefore, Dynamic *µ*-PBWT consumes approximately 1.84 bytes per site per haplotype for 1KGP data and approximately 1.5 bytes per site per haplotype for UKB data.

### 4.4 Long match query

We also tested the long match query algorithm on chromosome 21 of UK Biobank data. For this experiment, we queried 400 haplotypes against reference panels with 1000, 5000, 10000, 50000 and 100000 haplotypes for a length cutoff of 2000 sites. Table 1 shows the average query time per haplotype for different reference panel sizes. We observe that the query time per haplotype on Dynamic *µ*-PBWT and *µ*-PBWT is comparable. We also observe that the query time per haplotype increases sublinearly with the size of the panel.

**Table 1.** Long match query time per query haplotype.

| Panel Size | Dynamic μ-PBWT | | μ-PBWT | |
| --- | --- | --- | --- | --- |
|  | Avg. Time (in sec) | Std. Dev | Avg. Time (in sec) | Std. dev |
| 1000 | 0.029470 | 0.000840 | 0.026313 | 0.004631 |
| 5000 | 0.089743 | 0.003145 | 0.088879 | 0.005594 |
| 10000 | 0.125642 | 0.004762 | 0.160786 | 0.012228 |
| 50000 | 0.208166 | 0.010068 | 0.598709 | 0.067909 |
| 100000 | 0.232464 | 0.013217 | 1.077350 | 0.146789 |

## 5 Discussion

In this work, we introduced Dynamic *µ*-PBWT (which can also be seen as compressed d-PBWT), a dynamic and space-efficient variation of the PBWT data structure for fast haplotype matching. We used run-length compressed PBWT to achieve better compression rate and stored the runs in the B+ trees to enable dynamic updates. We developed algorithms to insert or delete a haplotype into/from the the Dynamic *µ*-PBWT without decompressing it. The number of updates per site per insertion or deletion in the B+ trees is constant regardless of the number of haplotypes in the Dynamic *µ*-PBWT. We showed that Dynamic *µ*-PBWT uses significantly less memory than d-PBWT. For example, for insertion, while Dynamic *µ*-PBWT is 60 times slower than d-PBWT, it consumes 37 times less memory than d-PBWT on a subsetted panel of UK Biobank data with 50000 haplotypes making it a space efficient choice for dynamic updates at biobank scale. We also provided a long match query algorithm on Dynamic *µ*-PBWT (missing in *µ*-PBWT) and showed that this algorithm can easily be extended back to the original static *µ*-PBWT.

While our algorithms are designed to be efficient, there is room for improvement in our current implementation. Dynamic *µ*-PBWT currently consumes roughly 33 times more memory than its static compressed counterparts. One way we could improve the memory consumption is to maintain a single B+ tree storing run length information of all the PBWT columns. This should improve our current compression rates as the overhead of maintaining multiple trees per column is reduced. Also, we used the DYNAMIC library in our current implementation of B+ trees but we believe a better implementation more specific to our use and appropriate for genetic data is possible. These improvements should make the compression rates comparable to static compressed PBWT and maintain similar or better insertion and deletion times.

The memory efficiency and flexibility of Dynamic *µ*-PBWT makes it a possible candidate data structure for biobank-scale genetic data. In response to the urgent need for efficient indexing and storage of large population data, Durbin noted PBWT as an efficient way to store haplotype data. Similarly, Li [10] mentions PBWT’s potential to be the primary way to share genetic data on which we can easily run analytic tasks. With the added functionalities of compression and dynamic updates above and beyond the original PBWT, we believe Dynamic *µ*-PBWT could be a potential common data format for biobank-scale genetic analysis ecosystem. Of course, further works are warranted to develop algorithms, codes, and use cases to cover all needed genetic analyses.

## Acknowledgement

This work was supported by the National Institutes of Health under award numbers R01HG010086 and R01AG081398. This research has been conducted using the UK Biobank Resource under Application Number 24247.

## Availability

The source code for Dynamic *µ*-PBWT is available at https://github.com/ucfcbb/Dynamic-mu-PBWT.

## Appendix

The following algorithms explain the FL mapping, insertion and deletion in Dynamic *µ*-PBWT and long match query on Dynamic *µ*-PBWT. Algorithm 2 uses Algorithm 1 as a subroutine to map a haplotype from column *k* to column *k* + 1. Algorithms 3, 4 and 8 are used as subroutines in the insertion algorithms defined in Algorithms 6 and 7. Algorithm 5 virtually inserts the new haplotype and calculates the divergence values of the new haplotype and the haplotype below it if the new haplotype were in the Dynamic *µ*-PBWT. Algorithms 6 and 7 insert the new haplotype into the Dynamic *µ*-PBWT one column at a time. Algorithms 9 and 10 delete the haplotype at a single column. For the long match query, Algorithm 5 is used to virtually insert the query haplotype *z* and Algorithm 11 finds all the long matches between query haplotype *z* and the haplotypes in Dynamic *µ*-PBWT.

### Algorithm 1

Algorithm to compute number of zeros, *u*, and number of ones, *v*, before *i* in a column with root node *x*

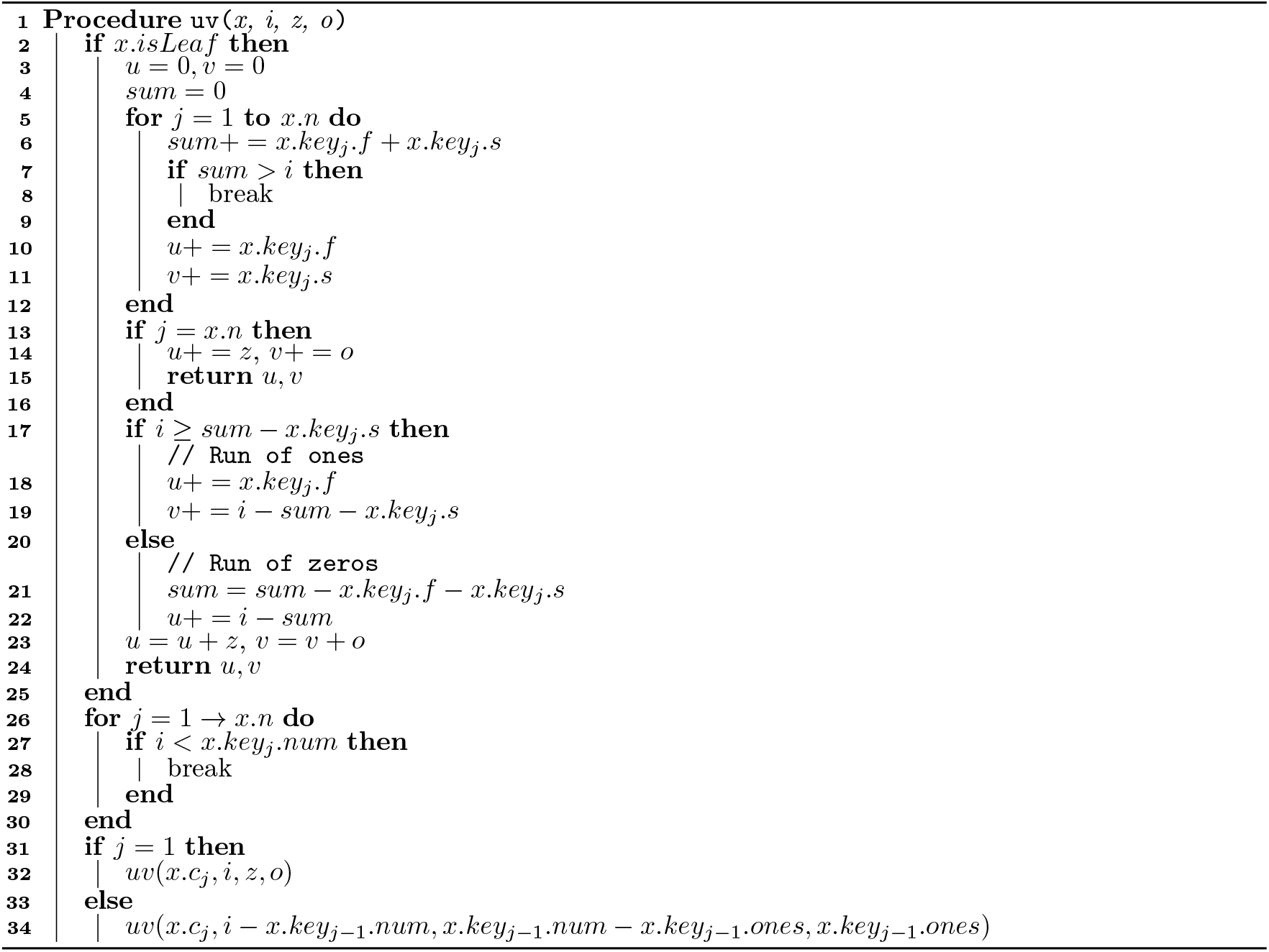

### Algorithm 2

Algorithm to map haplotype *id* with bit value *v* at *i*^*th*^ index in column *k* to column *k* + 1 using the tree of column *k* rooted at *x*

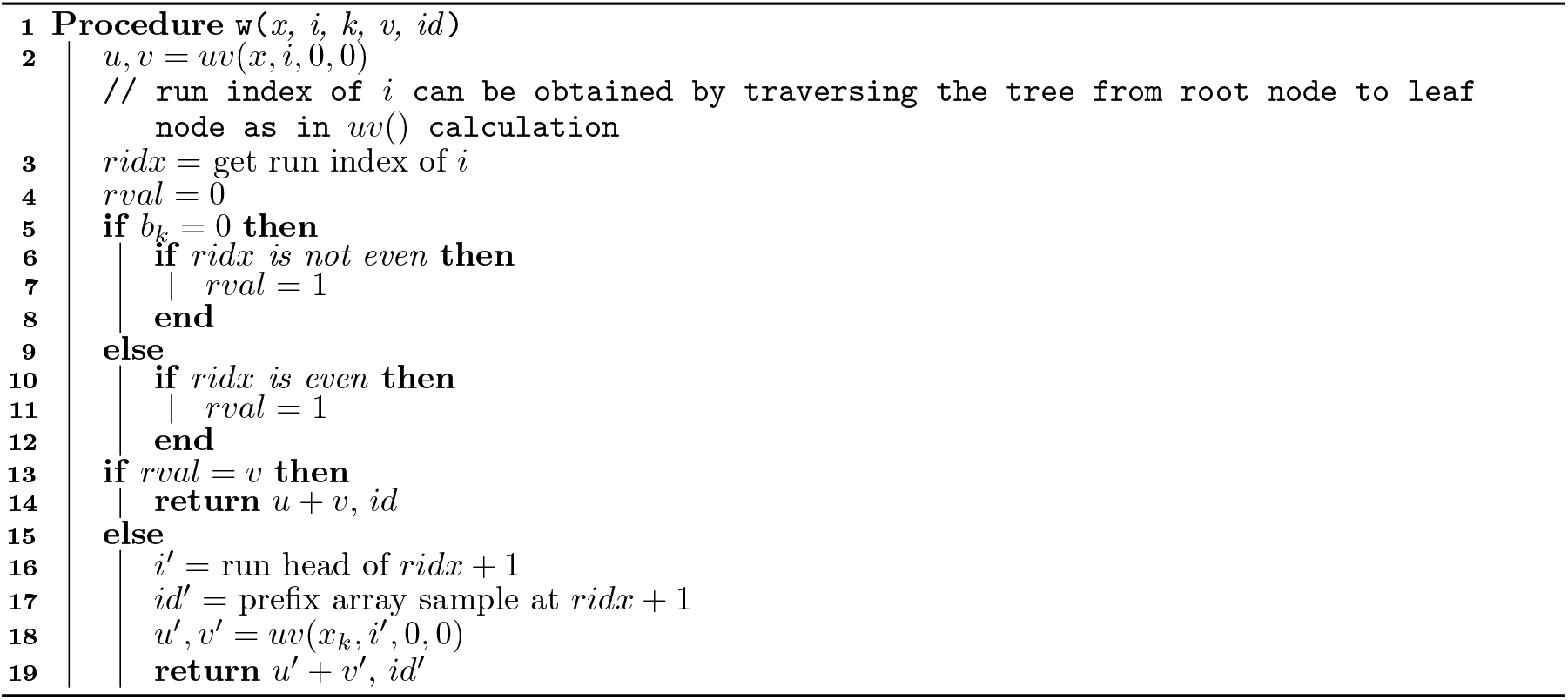

### Algorithm 3

Splitting algorithm for full leaf node *x.c*_*i*_ with non full parent node *x*

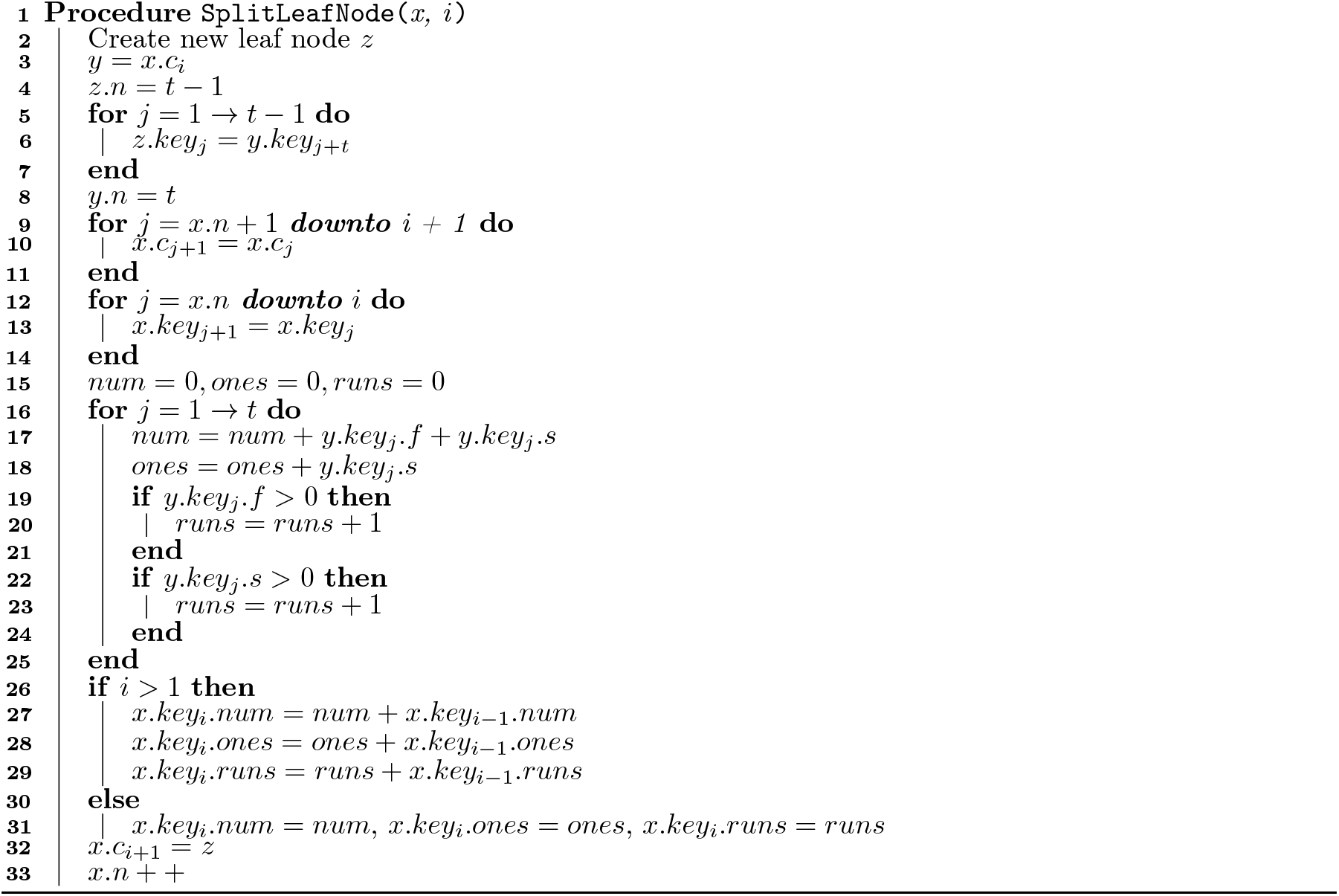

### Algorithm 4

Algorithm for splitting full internal node *c*_*i*_ with non-full parent node *x*

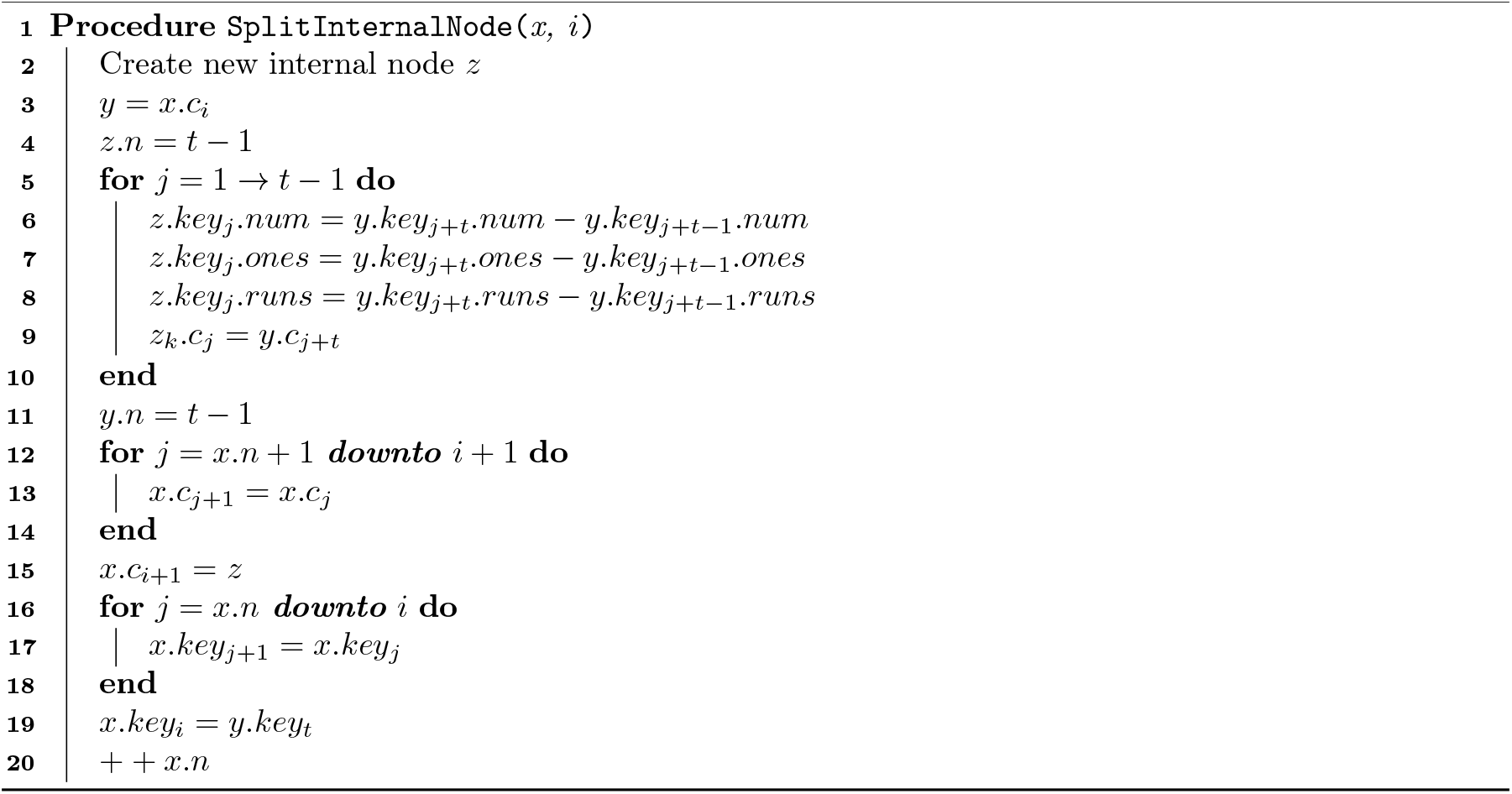

### Algorithm 5

Algorithm to virtually insert the query haplotype *z* and calculate the divergence values of *z* and the sequence below it if *z* were in the Dynamic *µ*-PBWT

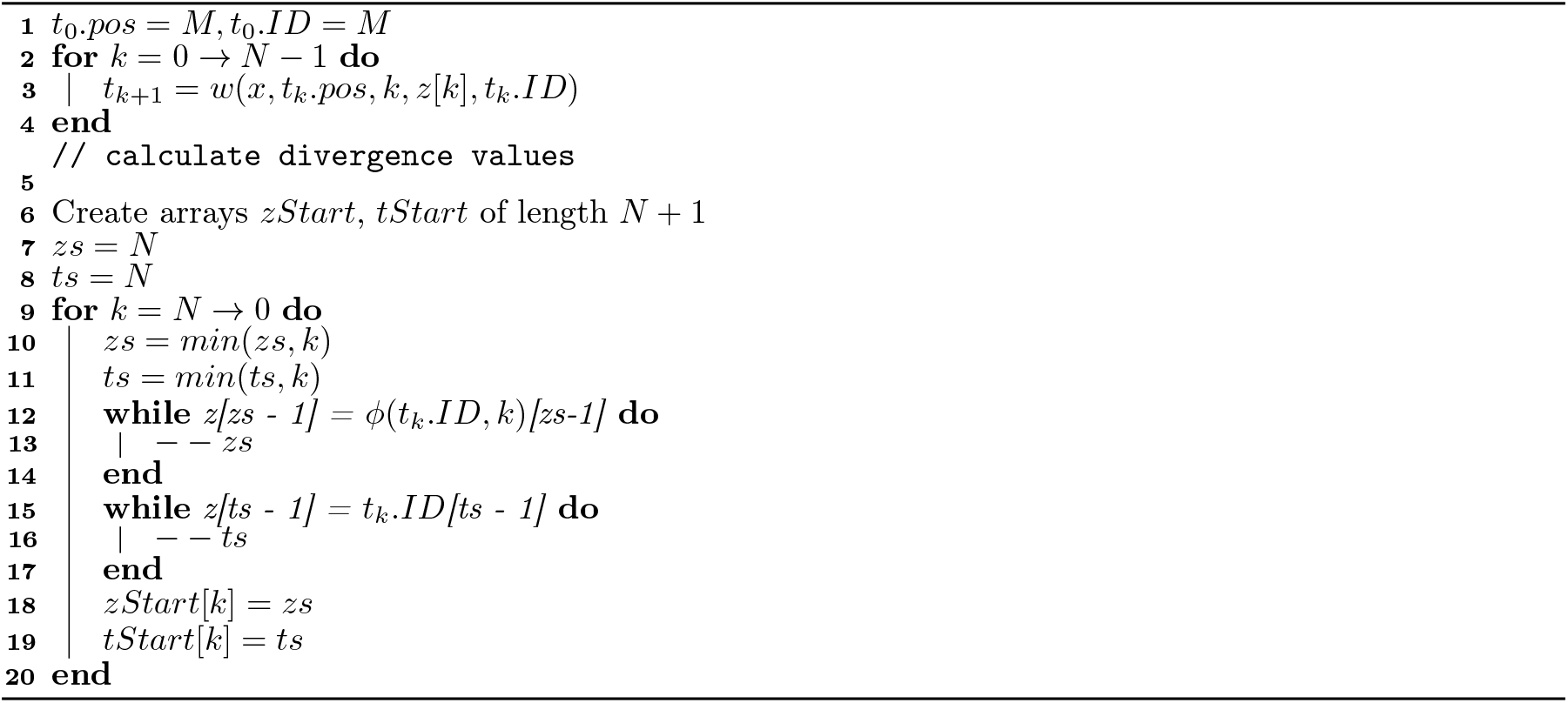

### Algorithm 6

Insertion algorithm for inserting a 0 in a tree at index *i* with root node *x*

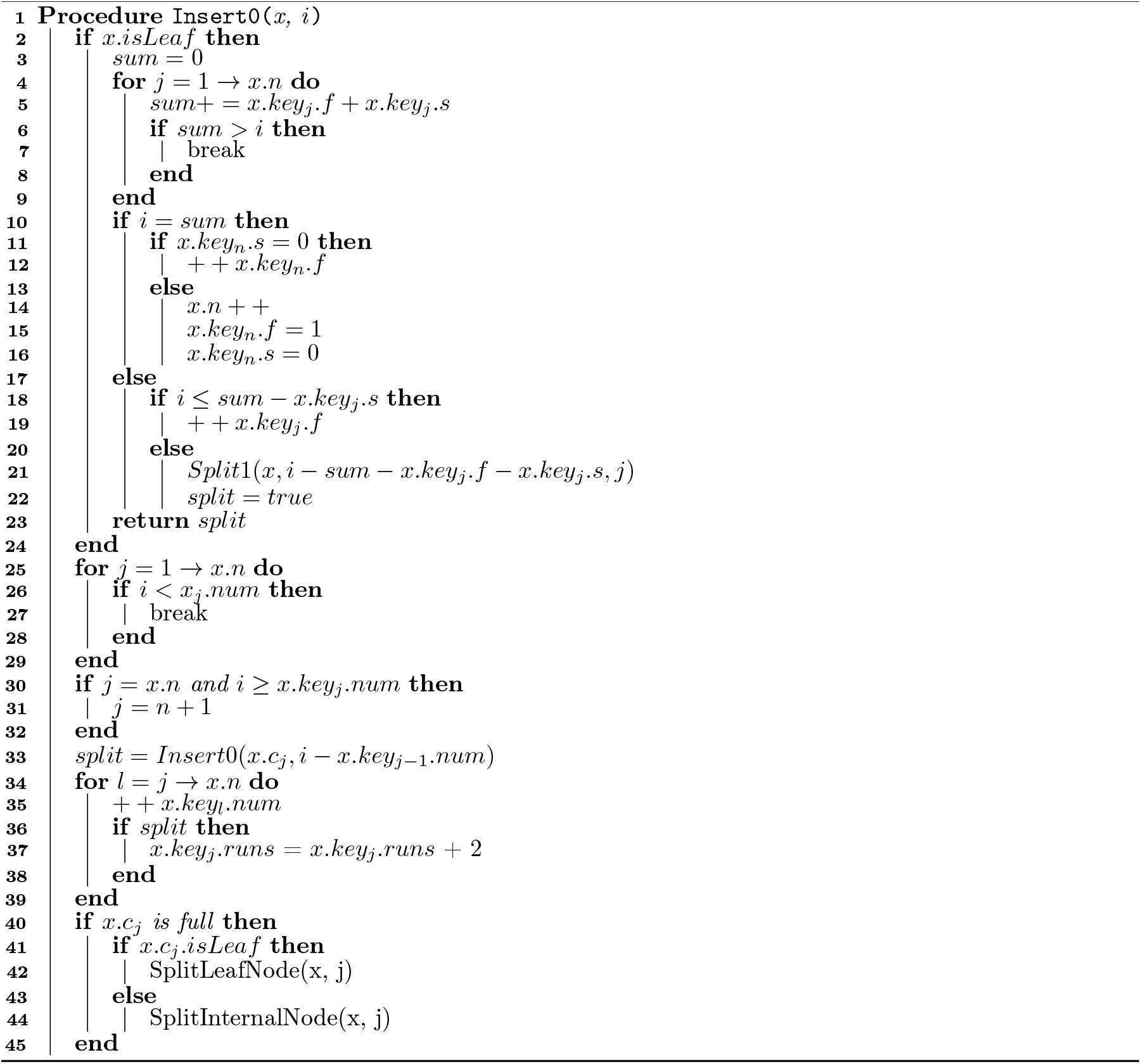

### Algorithm 7

Insertion algorithm for inserting 1 in a tree at index *i* with root node *x*

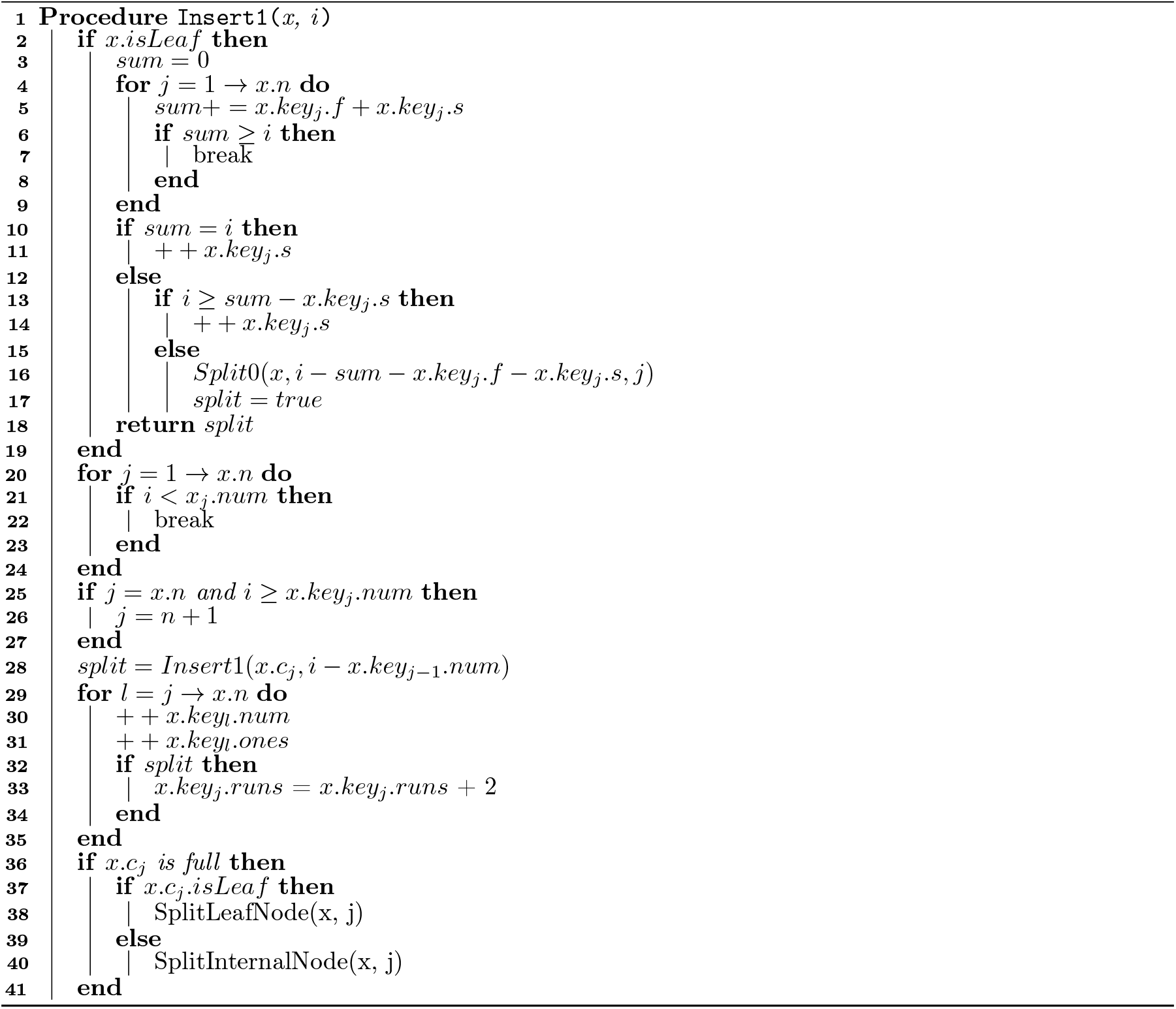

### Algorithm 8

Algorithm to split a run in the key *s* of a leaf node *x* where *leftHalf* is the number of bits in the run after splitting

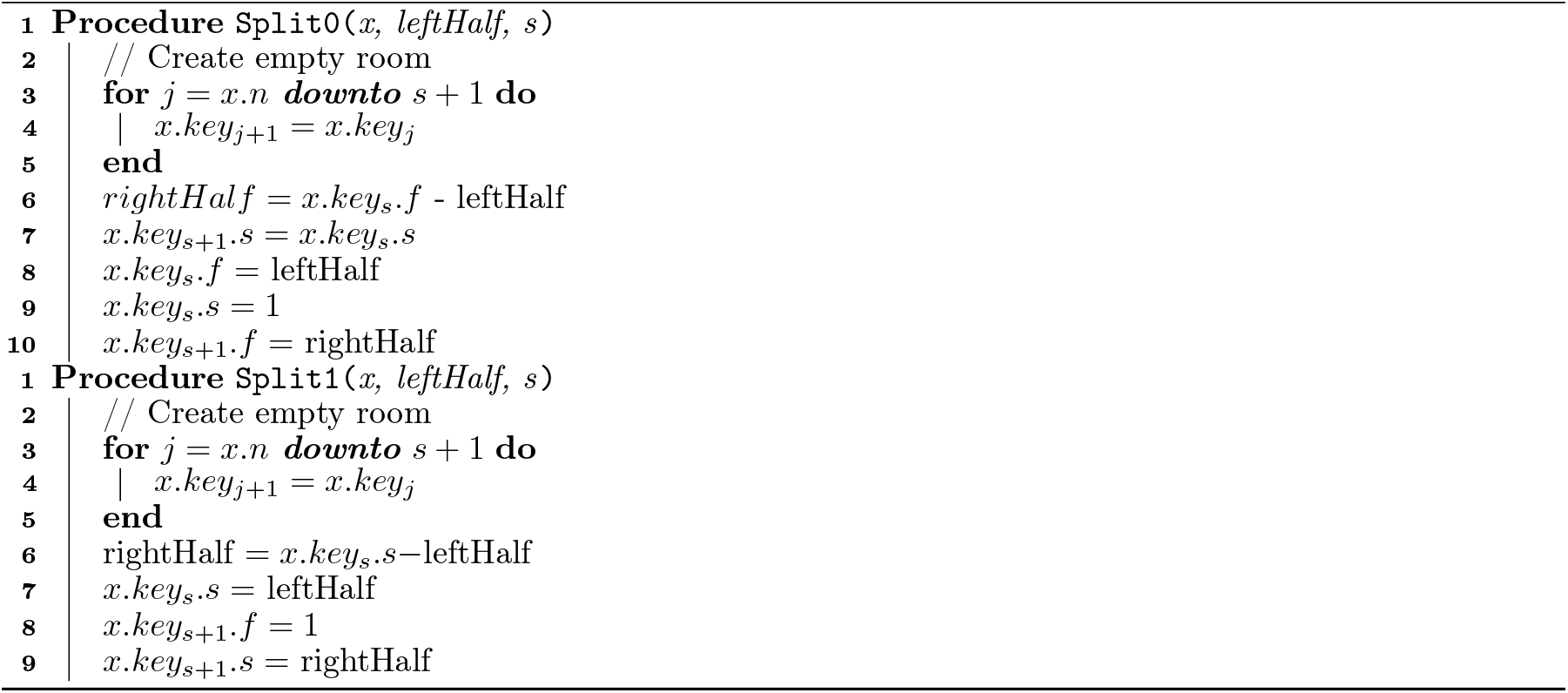

### Algorithm 9

Deletion Algorithm: Algorithm to delete 0 at index *i* in a tree with root node *x*

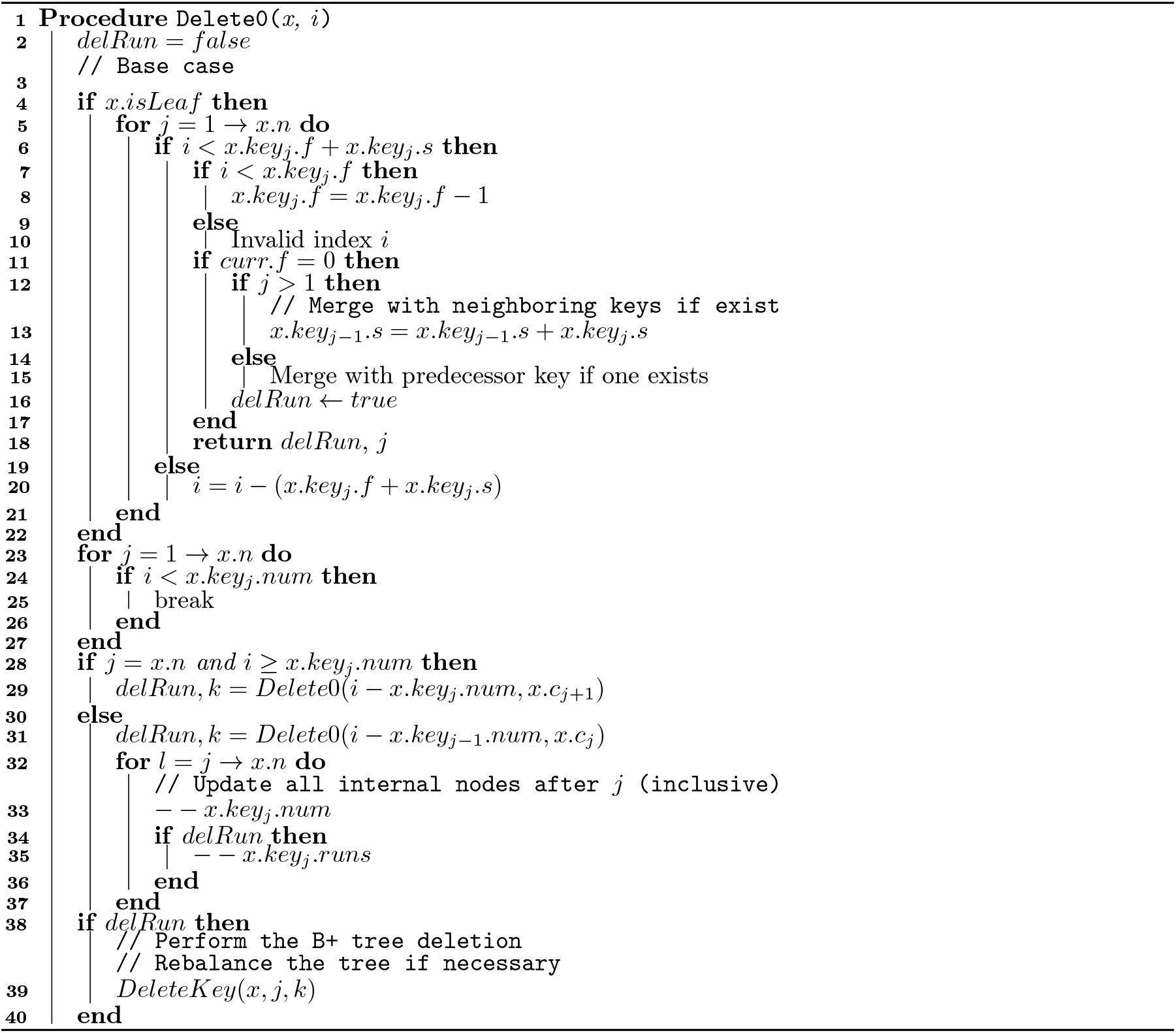

### Algorithm 10

Deletion Algorithm: Algorithm to delete 1 at index *i* in a tree with root node *x*

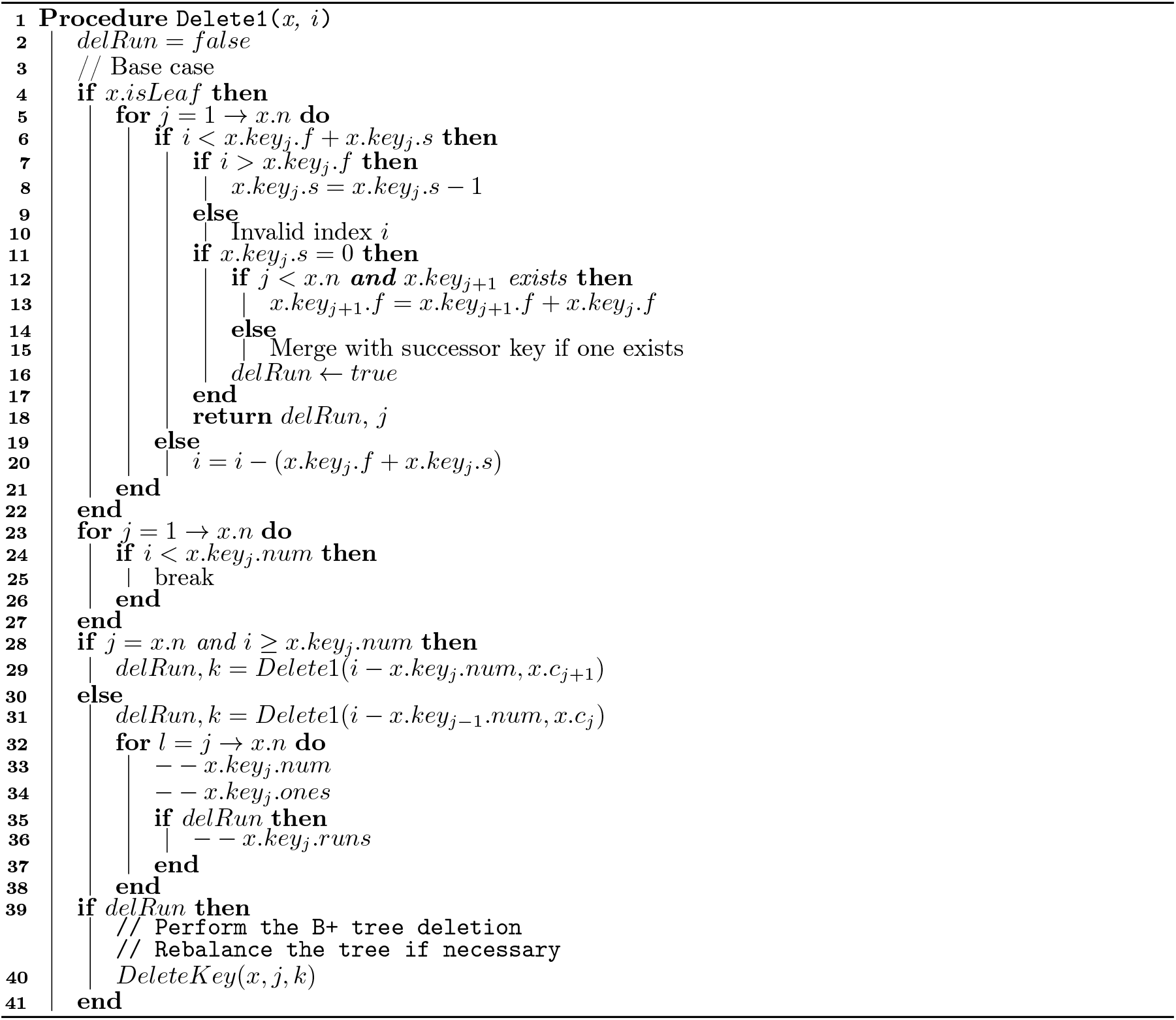

### Algorithm 11

Long match query algorithm to find matches at least length *L* between query haplotype *z* and sequences in Dynamic *µ*-PBWT

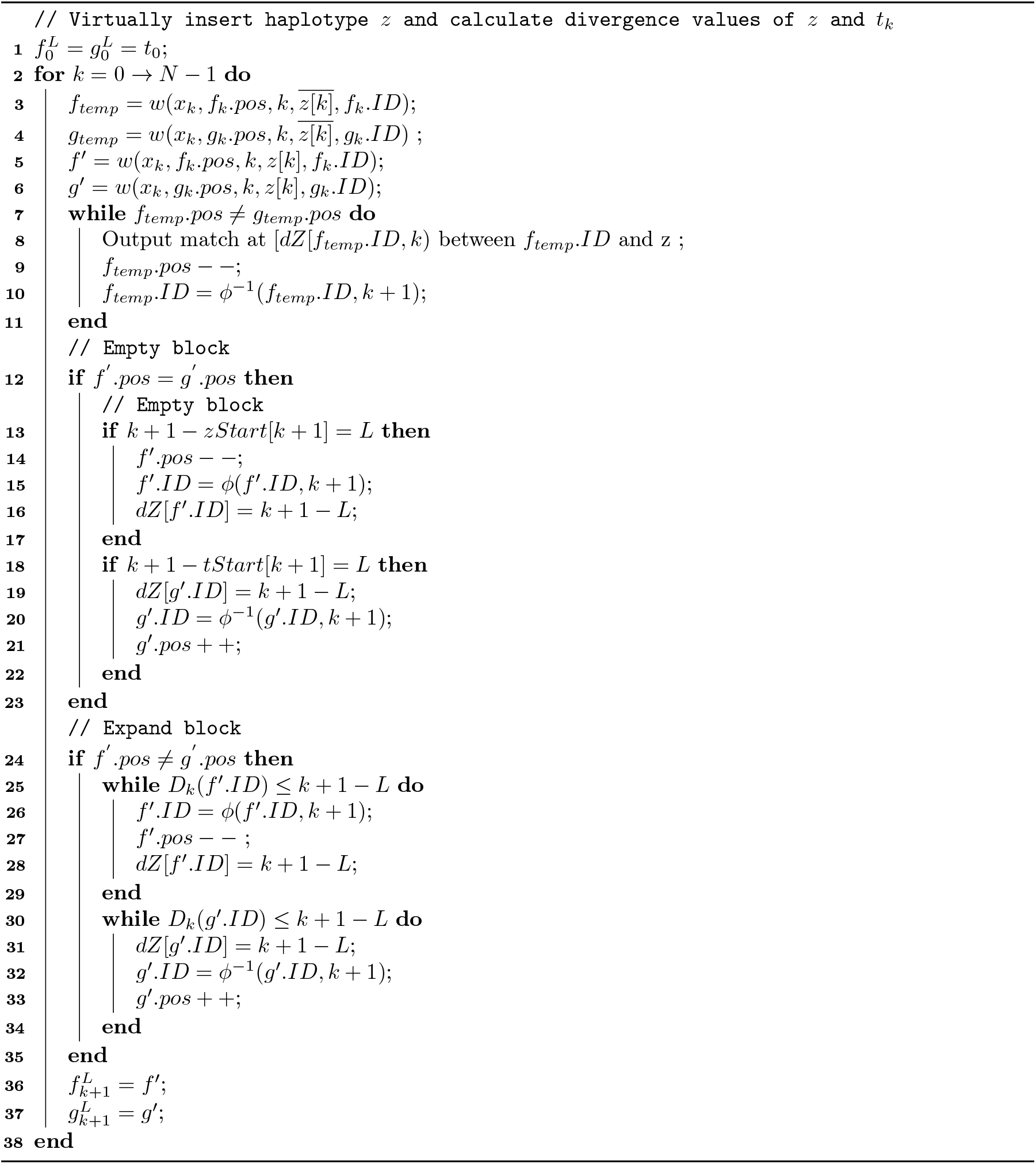

